# Differentiation-independent Activation of HPV Genome Replication by the lncRNA DINO

**DOI:** 10.64898/2026.06.19.733329

**Authors:** Warda Arman, Alison A. McBride, Karl Munger

**Author notes:** To whom correspondence should be addressed: Department of Developmental, Molecular and Chemical Biology, Tufts University School of Medicine, 136 Harrison Ave., MV701G Boston, MA 02111, USA.

## Abstract

Human papillomaviruses (HPV) rely on multiple host cell factors to replicate the viral genome, yet the contribution of host long noncoding RNAs (lncRNAs) to viral genome maintenance and amplification in the productive life cycle remains poorly understood. In this study, we show that the lncRNA DINO is a driver of HPV DNA replication. DINO levels increase during keratinocyte differentiation and ectopic expression of DINO promotes both HPV genome replication and the formation of replication foci, and this is independent of keratinocyte differentiation signals. Ectopic DINO expression increases select early viral transcript levels including E1^E4, E1, and E2. Notably, DINO’s subcellular localization is also context-dependent: during DNA damage DINO is predominantly cytoplasmic, but during keratinocyte differentiation nuclear retention is observed. This differential localization suggests that DINO has distinct functional roles in keratinocyte differentiation and HPV biology. Our findings highlight DINO as a lncRNA that promotes HPV genome replication and suggest that lncRNAs may play underappreciated roles in host-virus interactions. This work provides a foundation for further exploration of lncRNAs as potential therapeutic targets in HPV-associated diseases.

**Importance:** Human papillomaviruses (HPVs) are the causative agents of many anogenital tract and oral cancers, yet the host factors that trigger and support viral genome replication during the productive life cycle are incompletely understood. This study identifies the long noncoding RNA DINO as a host regulator that promotes HPV DNA replication, replication focus formation, and early viral gene expression independently of keratinocyte differentiation. We further show that DINO exhibits context-dependent subcellular localization, suggesting distinct functional roles in cellular stress responses and HPV biology. These findings reveal an underappreciated role for host lncRNAs in virus–host interactions and provide new insight into cellular pathways that support HPV genome replication.

## Introduction

Human papillomaviruses (HPVs) are the causative agents of approximately 5% of all cancers worldwide, including anogenital and head and neck carcinomas [1]. HPVs are categorized into high-risk and low-risk types based on their oncogenic potential. The high-risk types, HPV16 and HPV18 account for the majority of HPV-associated cancers, whereas HPV31 and other less prevalent high-risk types contribute to approximately 17% of cases [2].

HPVs are non-enveloped viruses with ∼8 kb double-stranded DNA genomes. The viral E1 and E2 proteins are essential for viral replication and transcription [3]. E5, E6 and E7 are viral accessory proteins, that manipulate host cell signaling pathways to promote long-term infections and enable viral genome replication in growth-arrested and terminally differentiated epithelial cells [4, 5]. The E4 protein is translated from a spliced E1^E4 mRNA and plays a crucial role in promoting genome amplification and viral synthesis [6]. The late proteins, L1 and L2, are the major and minor capsid proteins, respectively. L1 is sufficient to assemble into virus-like particles which are the basis of the current HPV vaccines. These vaccines have reduced the incidence of HPV-related lesions and cancers in vaccinated populations [7–9].

The HPV viral lifecycle is intricately tied to the host keratinocyte differentiation program [10–12]. HPVs infect actively dividing, undifferentiated basal keratinocytes, where viral genomes replicate and are maintained at a low copy number in the nuclei of infected cells [13]. During mitosis, HPV genomes are tethered to the mitotic chromosomes and are partitioned into daughter cells [14].

Papillomavirus genomes are maintained at a low copy number presumably as a strategy to evade immune detection in proliferative cells [15]. Mechanisms underlying papillomavirus genome maintenance have been studied extensively, and multiple models have been proposed. The traditional model posits that papillomavirus genome replication in proliferating keratinocytes is limited by viral repressors, such as the E8^E2 protein, which interacts with the nuclear receptor corepressor/silencing mediator for retinoid and thyroid hormone receptors (NCOR/SMRT) complex [16, 17] as well as host restriction factors such as Sp100 and IFI16 [18]. During the maintenance phase, the HPV genome may be replicated via two modes: an “ordered mode” where each genome is replicated in synchrony with host cell DNA synthesis, or a random mode where some genomes are replicated asynchronously whereas others are not. Overall, however, papillomavirus genomes are maintained at a consistent, low number [19]. HPV maintenance genome replication is thought to employ a bidirectional theta replication mechanism [20]. More recently, an alternative model was proposed, whereby HPV genomes are replicated at high numbers during the S and G2 phases of the cell cycle in basal epithelial cells. During mitosis, only a small number of genomes are tethered to mitotic chromosomes and the remaining, untethered viral genomes are not partitioned to daughter cells, and are degraded in the cytosol, thereby resetting the viral genome copy number after each cell division [21]. These contrasting mechanisms highlight the complexity of papillomavirus genome maintenance and the need for further investigation to delineate the precise pathways involved.

The HPV E6 and E7 proteins remodel host cellular pathways, shaping the cellular environment to support a persistent infection and a replication-competent state regardless of the specific mechanism of viral DNA replication [22, 23]. Viral genome amplification occurs in differentiated keratinocytes. High-level viral genome synthesis occurs in discrete nuclear compartments termed replication foci in a subset of differentiated cells [24–26]. The mechanism of HPV genome replication in the differentiated cells is thought to be by recombination-dependent replication [20, 27, 28]. The late viral transcripts, E1^E4 and later L1 and L2, are also expressed in these cells. In the most terminally differentiated cells, viral packaging occurs, and viral progeny remain in the denucleated squames when they are sloughed off the epithelium. HPV genome replication has been linked to cellular DNA damage repair pathways. HPV maintenance genome replication is independent of the Ataxia-Telangiectasia Mutated (ATM) kinase pathway but dependent on the Ataxia Telangiectasia and Rad3-related (ATR) kinase, whereas HPV genome amplification is requires both ATM and ATR [24, 29]. Various DNA repair factors are recruited to the viral replication foci to facilitate the synthesis of the viral genomes [30].

Most of the mechanistic work on HPV replication and pathogenesis has focused on delineating functional and biochemical interactions of HPV proteins with host cellular proteins. It is now clear, however, that the majority of the human transcriptome (∼98%) is not translated into proteins [31]. LncRNAs are defined as transcripts of >200 nucleotides that have limited coding potential and have been increasingly recognized to serve important functions in various cellular processes [32, 33].

The damage-induced long noncoding RNA, (DINO) was identified as a p53-responsive lncRNA that binds and stabilizes p53 in response to DNA damage, thereby amplifying p53 signaling [34]. In previous work with cervical carcinoma cells that contain integrated, subgenomic HPV16 sequences, we observed that DINO expression was upregulated following doxorubicin treatment, a potent inducer of double-stranded DNA breaks. Importantly however, DINO levels increased even when ATM was inhibited, indicating that DINO functions upstream of p53 in the DNA damage signaling pathway [35]. Ectopic acute DINO expression was sufficient to cause hallmarks of DNA damage such as 53BP1 foci, as well as ATM and p53 activation in the absence of exogenous DNA damage [35]. Hence DINO is not simply a p53-regulated p53 regulator and functions as an active component of DNA damage signaling. Based on these findings, we hypothesized that DINO may modulate HPV genome replication.

In this study, we demonstrate that the lncRNA DINO was induced in response to either DNA damage or differentiation of patient-derived cervical epithelial cell lines harboring HPV episomes. Ectopic expression of DINO promoted the formation of γH2AX foci and enhanced HPV genome replication, as evidenced by an increase in replication foci and increased mRNA levels of select viral transcripts E1^E4, E1 and E2. Notably, while DINO was upregulated during keratinocyte differentiation, its expression alone does not induce canonical differentiation markers, showing that DINO can promote viral replication independent from inducing a full keratinocyte differentiation program. Together, these findings point to a role for DINO in amplifying cellular signals that trigger HPV genome replication.

## Materials and Methods

### Plasmids

The pBR 9E HPV31 plasmid (HindIII-linearized) was used to generate fluorescence in situ hybridization (FISH) probes and served as a standard for HPV31 quantitative PCR (qPCR) assays [36]. The pMA-RPPH1 plasmid, containing a 341 bp fragment of the human RNase P gene, was used to generate standard curves for cellular gene quantification. The pEFHPV-16W12E plasmid, containing the HPV16 genome, was used as a standard for HPV16 qPCR (gift from Paul Lambert, University of Wisconsin, Madison, WI) [37]. For inducible DINO expression, the full-length human DINO transcript (a gift from Howard Chang, Stanford University) [34] was cloned into the doxycycline-inducible lentiviral vector pLIX_403-puro (Addgene plasmid #41395; gift from David Root, Broad Institute, Cambridge MA), as previously described [35]. A matched pLIX-based plasmid expressing green fluorescent protein (GFP) under the same inducible promoter was used as a control (gift from James DeCaprio, Dana Farber Cancer Institute, Boston, MA) [38].

### Cell lines and cell culture

CIN612-9E [39] and W12e [25] cell lines were maintained on mitomycin-C-treated mouse fibroblast 3T3-J2 feeder cells in F-media [39]. Cells were transduced with either DINO-expressing or GFP-expressing lentiviral vectors [35] and selected using 1 mg/ml puromycin. Doxycycline hyclate (Sigma-Aldrich) was added to F-media for the timepoints indicated at a final concentration of 1 µg/ml. All experiments were performed within five passages post-selection/post-thawing to prevent enrichment of cells with integrated viral genomes.

For calcium-induced differentiation, cells were grown to confluence in F-media with mitomycin-C-treated 3T3-J2 feeder cells. The media was then replaced with KBM (Lonza) supplemented with bovine pituitary extract, hydrocortisone, and epidermal growth factor for 24 hours.

Subsequently, the medium was replaced with KBM containing 1.5 mM calcium chloride (CaCl₂) for 4–5 days to induce differentiation.

For methylcellulose-induced differentiation, CIN612-9E cells were collected at 80% confluency. A total of 2 × 10⁶ cells were trypsinized, resuspended in 1 ml F medium, and added dropwise to a 10-cm petri dish containing 25 ml of a 1.5% methylcellulose solution. The methylcellulose solution was prepared by hydrating autoclaved methylcellulose (4,000 cps; cat. no. M0512, MilliporeSigma) in two equal volumes of F medium at 60°C for 20 min followed by stirring at 4°C for a minimum of 3 hours until clear, and finally adding FBS to 5%.. After 48 hours at 37°C in a humidified 5% CO₂ incubator, cells were harvested, washed four times with PBS, and pelleted by centrifugation.

### DNA damage experiments

For doxorubicin studies, Doxorubicin Hydrochloride (Fisher Scientific) was added to F-media at a final concentration of 0.2 µg/ml for 24 hours.

### RNA isolation and quantitative PCR

After removing the feeders from keratinocyte cultures, total RNA was extracted from cells using the Quick-RNA Miniprep Kit (Zymo Research), according to the manufacturer’s instructions. One microgram of RNA was reverse transcribed using the QuantiTect Reverse Transcription Kit (Qiagen) following the manufacturer’s instructions. Quantitative PCR (qPCR) was performed in triplicate using PowerUp SYBR Green Master Mix (Thermo Fisher Scientific) and thermal cycling conditions recommended by the manufacturer. Relative transcript levels were calculated using the comparative threshold cycle (ΔΔCt) method, normalizing to the reference gene RPLP0, and are reported as fold change relative to the indicated control condition. Primer sequences are listed in Table 1.

**Table 1.**
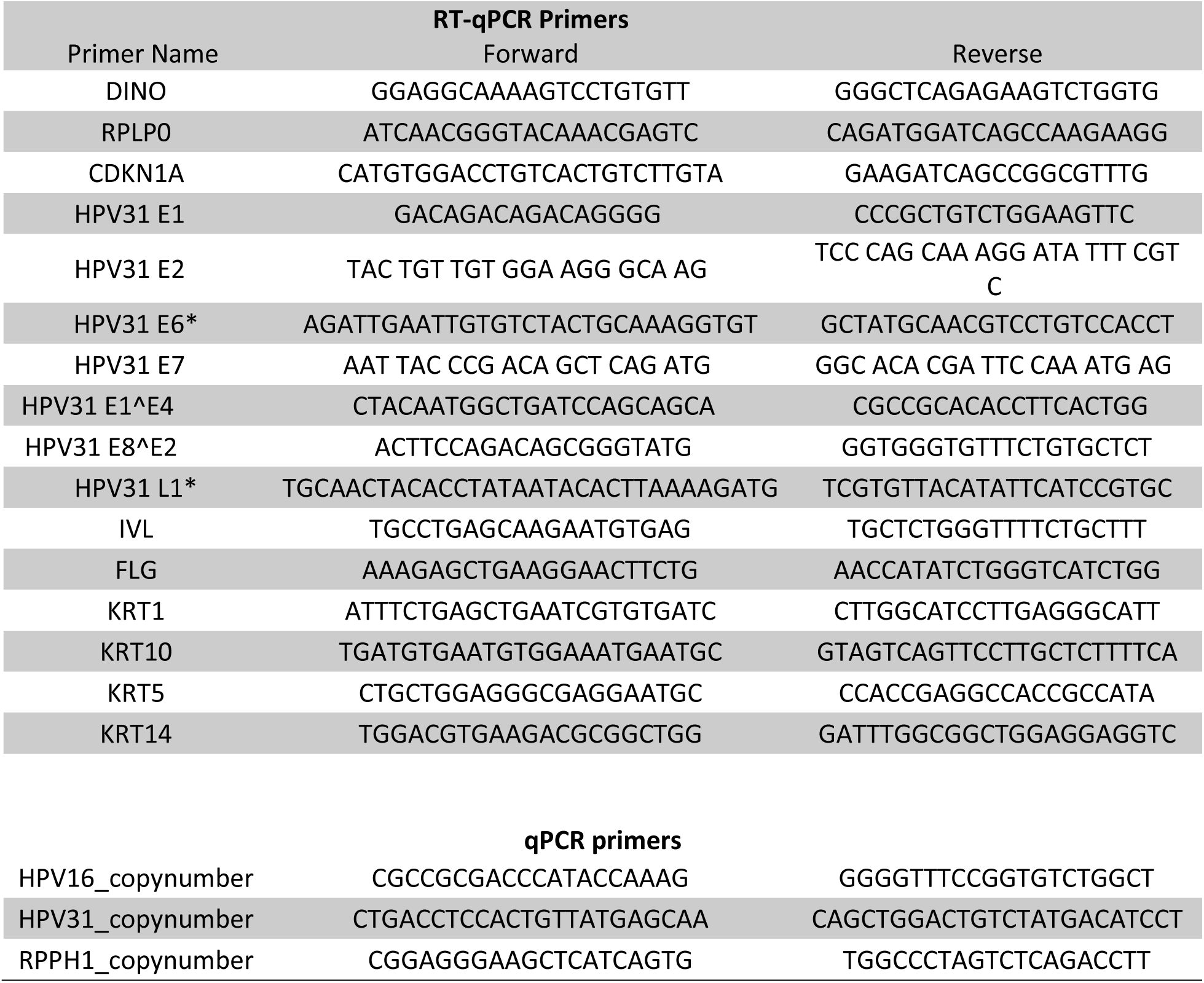
PCR primers.

### HPV DNA copy number

After removing the feeders from keratinocyte cultures, total cellular DNA was isolated from cells using the DNeasy Blood and Tissue kit (Qiagen). 40 ng of DNA was analyzed by qPCR using PowerUp SYBR Green Master Mix (Thermo Fisher Scientific) and thermal cycling conditions recommended by the manufacturer. HPV DNA copy number was determined by comparison to standard curves generated from serial dilutions of the pBR 9E HPV31 plasmid or the pEFHPV-16W12E plasmid, which contain the full-length HPV31 and HPV16 genomes, respectively. Primer sequences are listed in Table 1.Copy number of the ribonuclease P RNA component H1 gene (RPPH1) was determined using a standard curve generated from the pMA-RPPH1 plasmid. Data are presented as fold change relative to GFP-expressing cells at 0 hours or prolferative conditions in differentiation experiments.

### Western blotting and antibodies

Cells were lysed in radioimmunoprecipitation assay (RIPA) buffer supplemented with Pierce protease and phosphatase inhibitors (Thermo Scientific) at 4°C for 30 minutes. Lysates were cleared by centrifugation at 15,000 × g for 15 minutes, and equal amounts of protein (50 μg) were fractionated on 4–12% NuPAGE Bis-Tris gels (Invitrogen) and transferred to polyvinylidene difluoride (PVDF) membranes (Millipore). Membranes were blocked in TBS-T containing 5% nonfat dry milk for 1 hour at room temperature and incubated overnight at 4°C with primary antibodies against TP53 (1:1,000; OP43, Calbiochem) and actin (1:1,000; MAB1501, Sigma-Aldrich). Membranes were washed in TBS-T and incubated with HRP-conjugated anti-mouse secondary antibody (1:10,000; NA931V, Invitrogen) for 1 hour at room temperature. Bands were visualized by enhanced chemiluminescence and digitally acquired on a G:Box Chemi-XX6 imager with Genesys software (Syngene).

### Immunofluorescence

Cells were grown on coverslips, fixed in 4% paraformaldehyde in PBS for 15 minutes at room temperature, and permeabilized in 0.1% Triton X-100/PBS for 15 minutes. Non-specific binding was blocked with 5% normal donkey serum in PBS for 1 hour at room temperature. Cells were incubated with rabbit anti-phospho-histone H2A.X (Ser 139) (Cell Signaling; #9718; 1:500) for 1 hour at 37°C. Coverslips were washed three times in PBS for 5 minutes each and then incubated with Rhodamine Red™-X (RRX) AffiniPure® Donkey Anti-Rabbit secondary (Jackson ImmunoResearch Labs; # 711-295-152; 1:100) for 1 hour at 37°C. Nuclei were stained with DAPI (4’,6-diamidino-2-phenylindole) and coverslips mounted in ProLong Diamond Antifade Mountant (Invitrogen).

### Fluorescence in situ hybridization (FISH)

Cells were grown on coverslips and fixed in fresh cold methanol and acetic acid (3:1) at -20°C for 10 minutes, followed by drying overnight at room temperature and storage at -20°C. HPV31 DNA FISH probes were prepared using the FISH-Tag DNA Multicolor Labeling Kit (Life Technologies), according to the manufacturer’s protocol. Hybridization was performed overnight at 37°C in 1× hybridization buffer (Empire Genomics) with 75 ng of labeled probe DNA per coverslip. Slides were washed with 1× phosphate-buffered detergent (PBD; MP Biosciences) at room temperature, followed by washing with 0.5× SSC and 0.1% SDS at 65°C. Nuclei were stained with DAPI, and coverslips were mounted using ProLong Gold Antifade Mountant (Life Technologies). Images were captured using a TCS-SP8 confocal microscope (Leica Microsystems). ImageJ was used to count nuclei per image, and cells containing HPV31 FISH foci were manually scored to determine the percentage of cells with at least one focus. A focus was defined as any discrete, punctate FISH signal visible above background within the nucleus, regardless of size or intensity. Images were anonymized prior to scoring so that the investigator was blinded to experimental conditions. A minimum of 300 nuclei were scored per replicate and condition.

### RNA Scope

Cells were grown on coverslips, fixed in 4% paraformaldehyde in PBS for 30 minutes at room temperature, and dehydrated through a graded ethanol series (50%, 70%, and 100%) for 1 minute each. Coverslips were stored in 100% ethanol at -20°C. Hybridization with DINO-specific probes (RNAscope™ Probe-Hs-DINOL catalog number 124572; Advanced Cell Diagnostics) was conducted according to the manufacturer’s instructions for the RNAscope Multiplex Fluorescence v2 kit (catalog number 323100; Advanced Cell Diagnostics). Following the HRP (horseradish peroxidase) blocker and final wash steps, coverslips were mounted on glass slides using ProLong Diamond Antifade Mountant (Life Technologies).

### Image Analysis

Except for FISH scoring, all microscopy images were collected on a Zeiss LSM800 laser scanning confocal microscope equipped with a 63× Plan-Apochromat oil immersion objective (NA = 1.4). Images were acquired as 0.5 μm z-stacks. For HPV replication foci and DINO RNA Scope quantification, maximal projection intensity images were analyzed using the Aggrecount [40] plugin on ImageJ. “Aggregates”, which we used to identify foci and DINO RNA Scope dots, were defined as having a minimum size of 0.39 µm diameter.

## Results

### DINO levels increase in response to DNA damage in patient-derived, HPV episome-containing cervical epithelial cell lines

It was previously shown that DINO is induced in response to doxorubicin, a chemotherapy agent that potently induces double-strand DNA breaks, in HPV16 positive cervical carcinoma cell lines and in human fibroblasts [34, 35]. We sought to determine whether this effect is conserved in in patient-derived, high-risk HPV episome-containing cell lines where all the viral genes are expressed. We evaluated endogenous DINO expression in W12e (HPV16) [39, 41] and CIN-612 9E (HPV31) [25] cell lines, which support the differentiation-dependent HPV lifecycle. Cells were treated with doxorubicin and DINO expression was evaluated using quantitative reverse transcription polymerase chain reaction (qRT-PCR). As a readout for double-strand DNA breaks we evaluated γH2AX, the serine 139 (Ser139) phosphorylated form of histone H2AX, that marks sites of double-strand DNA breaks [42]. Immunostaining and quantification of γH2AX foci revealed a significant increase (84.73 foci per cell) in the number of γH2AX foci in doxorubicin-treated cells (Figure 1A), consistent with elevated DNA damage signaling in response to double-strand DNA breaks. DINO expression increased following doxorubicin treatment in both the CIN-612 9E (98.71 +/-31.61 fold) and W12e cell lines (6.20 +/- 1.13 fold) (Figure 1B). Hence, DNA damage triggers DINO expression in these cell lines.

**Figure 1.**
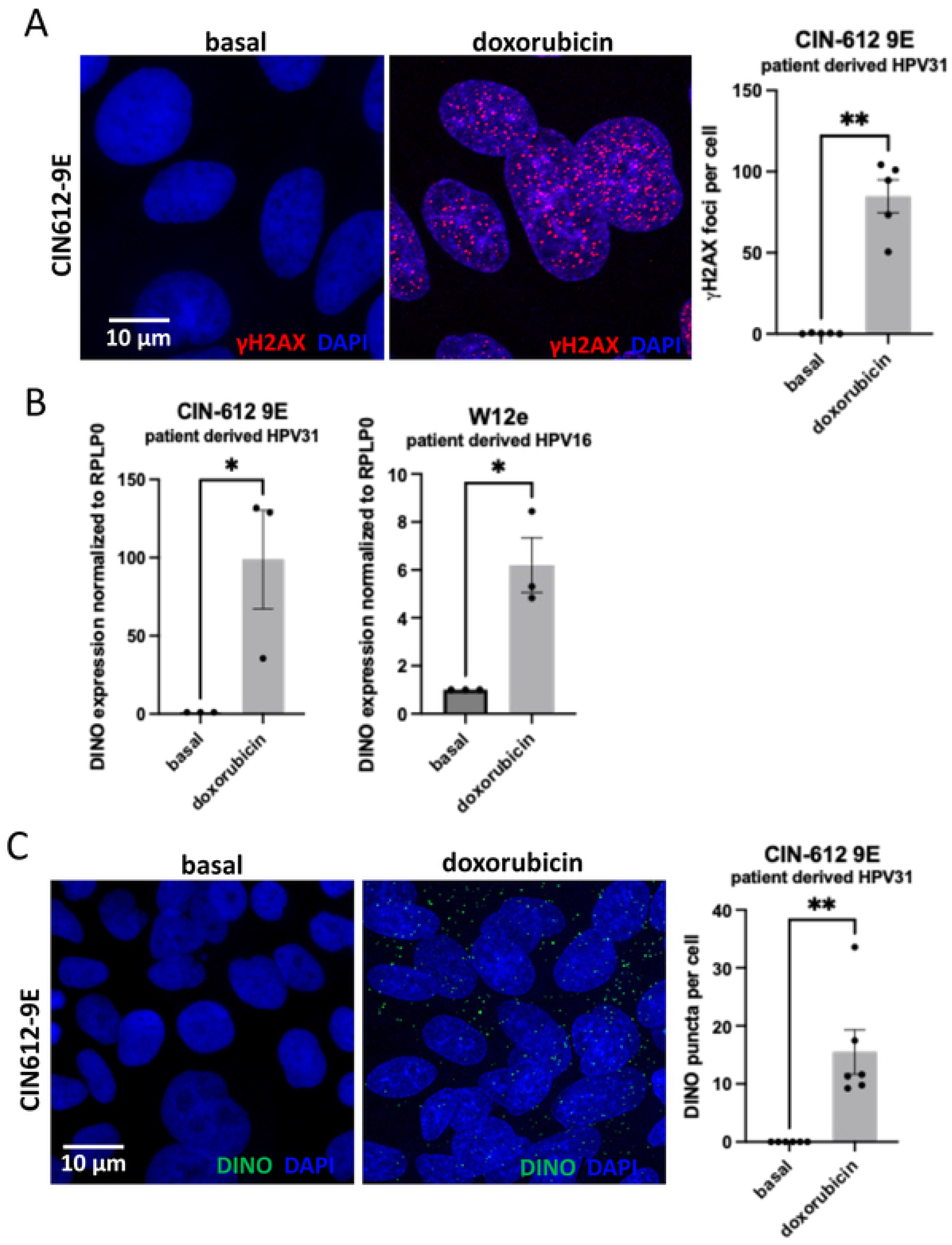
DINO levels increase in response to DNA Damage in patient-derived, HPV episome containing cervical epithelial cell lines. (A) Immunofluorescence microscopy of CIN612-9E cells treated with doxorubicin for 24 h, stained for γH2AX to detect DNA double-strand breaks. Quantification of γH2AX foci per nucleus confirms activation of the DNA damage response. Data are presented as mean ± SEM from one biological replicate with two technical replicates (two coverslips); each point represents one image containing 20 to 50 nuclei. The experiment was repeated twice with similar results. An unpaired t-test with Welch’s correction was used for statistical comparisons. (B) Quantitative PCR analysis of DINO expression in untreated (basal) and doxorubicin-treated CIN612-9E and W12 cells. DINO transcript levels were normalized to RPLP0. Data are presented as mean ± SEM from three biological replicates, each with two technical replicates. Statistical comparisons were performed using an unpaired t-test. (C) RNA scope analysis of DINO transcripts following doxorubicin treatment. Quantification of DINO puncta per cell shows an average increase of 15 transcripts. Data are presented as mean ± SEM from one biological replicate with two technical replicates (two coverslips); each point represents one image containing ∼25 nuclei. The experiment was repeated three times with similar results. An unpaired t-test with Welch’s correction was used for statistical comparisons.

To assess the copy number and subcellular localization of DINO transcripts, we performed RNA scope analysis in control and doxorubicin-treated CIN612-9E cells (Figure 1C). Quantification of DINO puncta, with each punctum representing a single transcript, revealed an average increase of **∼**15 transcripts (15.49) per cell following treatment. Notably, this value represents the mean across all cells, as individual cells displayed considerable variability in transcript abundance.

### DINO levels increase during differentiation of HPV episome-containing cervical epithelial cell lines

Given that DNA damage signaling is critical for viral genome replication [43–45] and that DNA damage signaling is important for keratinocyte differentiation [46], we next investigated whether DINO levels increase during keratinocyte differentiation. Keratinocyte differentiation was induced by adding 1.5 mM Ca^2+^ to keratinocyte serum-free medium, and DINO levels were assessed by qRT-PCR. To confirm that this procedure induced differentiation, the levels of the keratinocyte differentiation markers, involucrin and filaggrin, as well as expression of the late viral transcript L1 were assessed (Figure 2A). Upon differentiation we observed increases in both involucrin and filaggrin as well as L1 indicating that the cells have differentiated appropriately. DINO levels robustly increased in response to differentiation in both cell lines (Figure 2B). This effect was not specific to Ca^2+^-induced differentiation nor the presence of the HPV virus since DINO expression also increased when CIN612-9E cells were differentiated by suspension in methylcellulose or HPV negative human foreskin keratinocytes were differentiated (Supplementary Figure 1). We further employed RNA scope to examine the copy number and subcellular localization of DINO transcripts during differentiation, observing an average of ∼50 nuclear transcripts (51.03) in differentiated cells (Figure 2C) compared to less than one in proliferative cells. These findings demonstrate that DINO levels increase during the differentiation of high-risk HPV episome–containing cervical epithelial lines and it is predominantly nuclear compared to induction by DNA damage.

**Figure 2.**
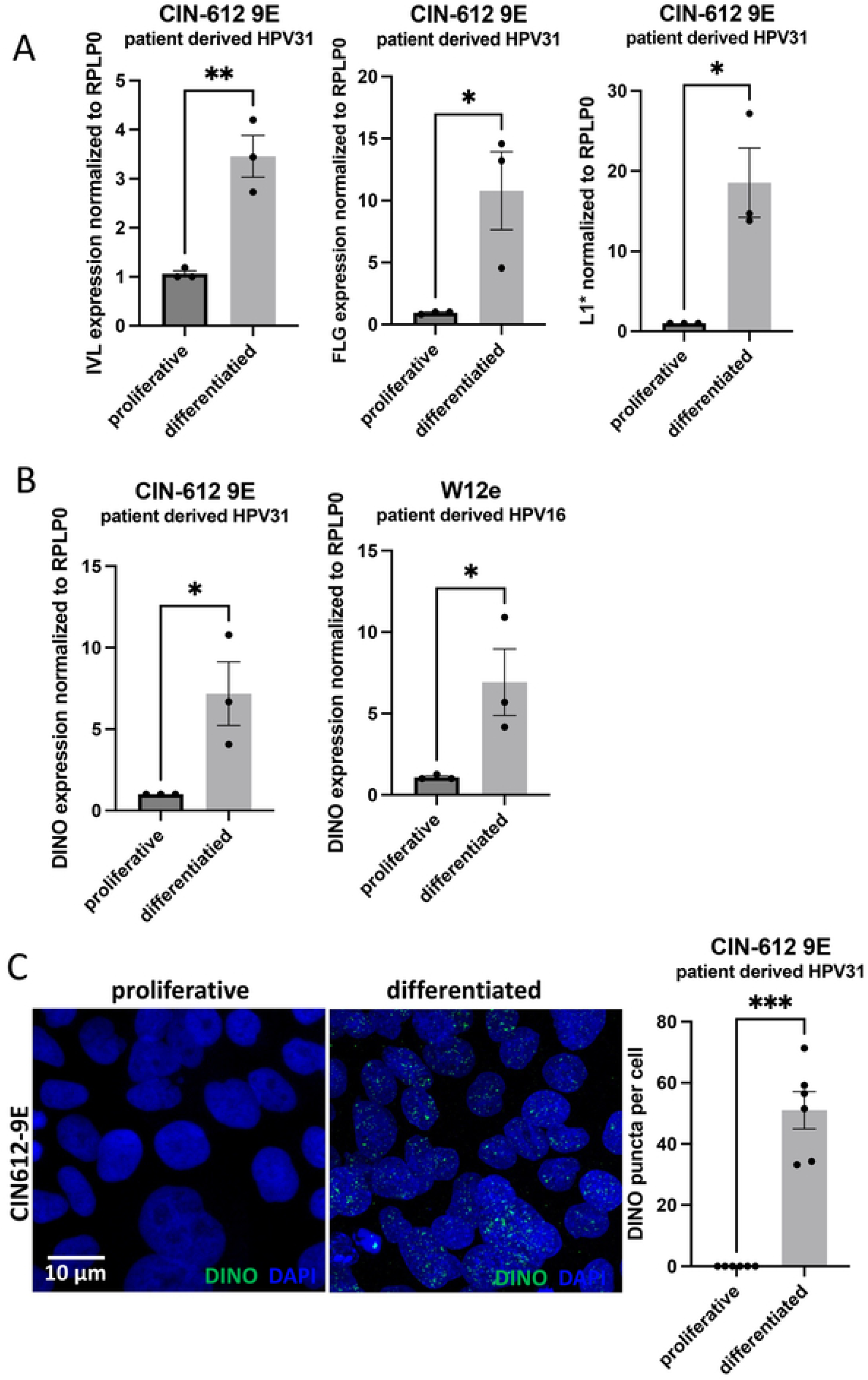
Differentiation induces DINO expression in HPV episome-containing cervical keratinocytes. (A) qPCR analysis of differentiation markers in CIN612-9E cells following induction of differentiation. Expression of IVL (involucrin), FLG (filaggrin), and the HPV late gene transcrip L1 increased, confirming successful induction of differentiation. Data are presented as mean ± SEM from three biological replicates, each with two technical replicates. Statistical comparisons were performed using an unpaired t-test. (B) DINO expression levels measured by qPCR in W12 and CIN612-9E cells before and after differentiation. Data are presented as mean ± SEM from three biological replicates, each with two technical replicates. Statistical comparisons were performed using an unpaired t-test. (C) RNA scope analysis of DINO transcripts following differentiation. Quantification of DINO puncta per nucleus shows an average of 50 transcripts per nucleus. Data are presented as mean ± SEM from one biological replicate with two technical replicates (two coverslips); each point represents one image containing ∼25 nuclei. The experiment was repeated three times with similar results. Statistical comparisons were performed using an unpaired t-test with Welch’s correction.

### Ectopic DINO expression promotes γH2AX focus formation in HPV episome-containing cells without detectable p53 activation

To explore the role of DINO in DNA damage signaling, we generated doxycycline-inducible DINO- or GFP-expressing W12e and CIN612-9E cell lines. Upon doxycycline treatment, DINO expression was robustly induced compared to doxycycline-treated GFP-expressing control cells (Figure 3A). We previously reported that ectopic DINO expression in the HPV16-positive SiHa and CaSki cervical carcinoma cells caused increased p53 tumor suppressor activity as assessed by increased expression of the canonical p53 transcriptional target gene *CDKN1A,* which encodes the p21^CIP1^ cyclin-dependent kinase inhibitor [35]. However, these lines express E6 and E7 from integrated HPV16 sub genomes [47]. To determine if ectopic DINO expression in CIN612-9E and W12e cells stabilized p53, we conducted immunoblot analysis. Steady state levels of p53 remained unchanged after 48 hours of DINO expression, whereas GFP expression caused a modest decrease in p53 levels (Figure 3B). To confirm that there was no increase in p53 transcriptional activity, we analyzed *CDKN1A* mRNA expression levels. In contrast to what we previously observed in SiHa and CaSki cells [35], DINO expression did not cause an increase in *CDKN1A* mRNA expression in the two HPV episome-containing cell lines (Figure 3C). However, ectopic DINO expression is sufficient to induce hallmarks of DNA damage signaling, as evidenced by an increase in γH2AX foci in CIN612-9E cells (Figure 3D). Hence, DINO expression is sufficient to trigger DNA damage signaling in HPV episome-containing cells, without detectable p53 activation.

**Figure 3.**
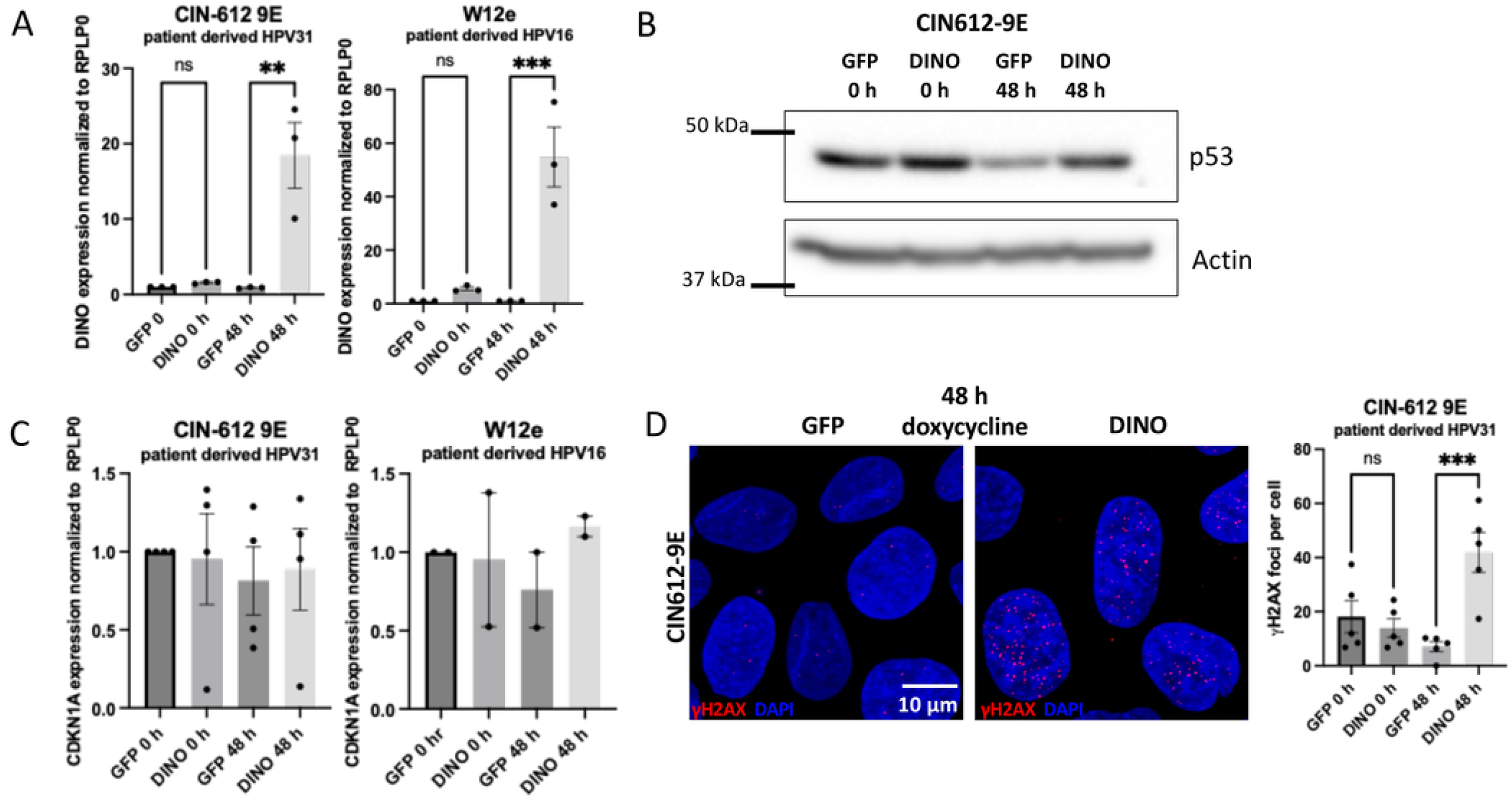
Ectopic DINO expression promotes γH2AX focus formation in HPV episome-containing cells independently of p53 stabilization. (A) CIN612-9E and W12e cells were transduced with doxycycline-inducible GFP or DINO vectors and treated with doxycycline for 48 hours. qRT-PCR confirmed DINO induction in the DINO-expressing cells, while transcript levels remained unchanged in the GFP controls. Data are presented as mean ± SEM from three biological replicates, each with two technical replicates. Statistical comparisons were performed using one-way ANOVA. (B) Western blot analysis of p53 protein levels following 48 hours of doxycycline treatment of GFP and DINO-expressing CIN612-9E cells. p53 steady state levels were unchanged upon ectopic DINO expression. (C)) CDKN1A expression levels were quantified by qPCR following 48 hours of doxycycline treatment and transcript levels remained unchanged in both CIN612-9E and W12e cells. (D) Immunofluorescence staining for γH2AX in CIN612-9E cells after 48 hours of doxycycline treatment. Quantification of γH2AX foci per nucleus showed a significant increase in DINO-expressing cells compared to matched GFP controls. Data are presented as mean ± SEM from one biological replicate with two technical replicates (two coverslips); each point represents one image containing ∼25 nuclei. The experiment was repeated three times with similar results. An unpaired t-test with Welch’s correction was used for statistical comparisons.

### Nuclear DINO transcripts are enriched during differentiation in both HPV-containing cells and primary keratinocytes

Next, we employed RNA scope to quantify DINO transcripts per cell in this inducible system. Following doxycycline treatment, DINO-expressing cells contained an average of ∼27 transcripts per cell (26.86) in CIN612-9E cells (Figure 4A), comparable to the levels observed after doxorubicin treatment. Analysis of DINO localization revealed a distinct pattern: during doxorubicin treatment, the majority of DINO transcripts were cytoplasmic (71.1%), whereas differentiation induced nuclear enrichment of transcripts (70.6%). Ectopic DINO expression produced an intermediate phenotype, with transcripts distributed between the cytoplasm (46.3%) and nucleus (53.7%) (Figure 4B, last panel). To investigate whether a primarily nuclear DINO localization pattern upon differentiation was also observed in HPV negative keratinocytes, we differentiated primary human foreskin keratinocytes and conducted RNA scope analysis. As expected, the number of DINO transcripts increased during differentiation to ∼22 transcripts per cell (21.97) (Figure 4C) and the majority of transcripts were nuclear in differentiated HFKs (69.6%) (Figure 4D).

**Figure 4.**
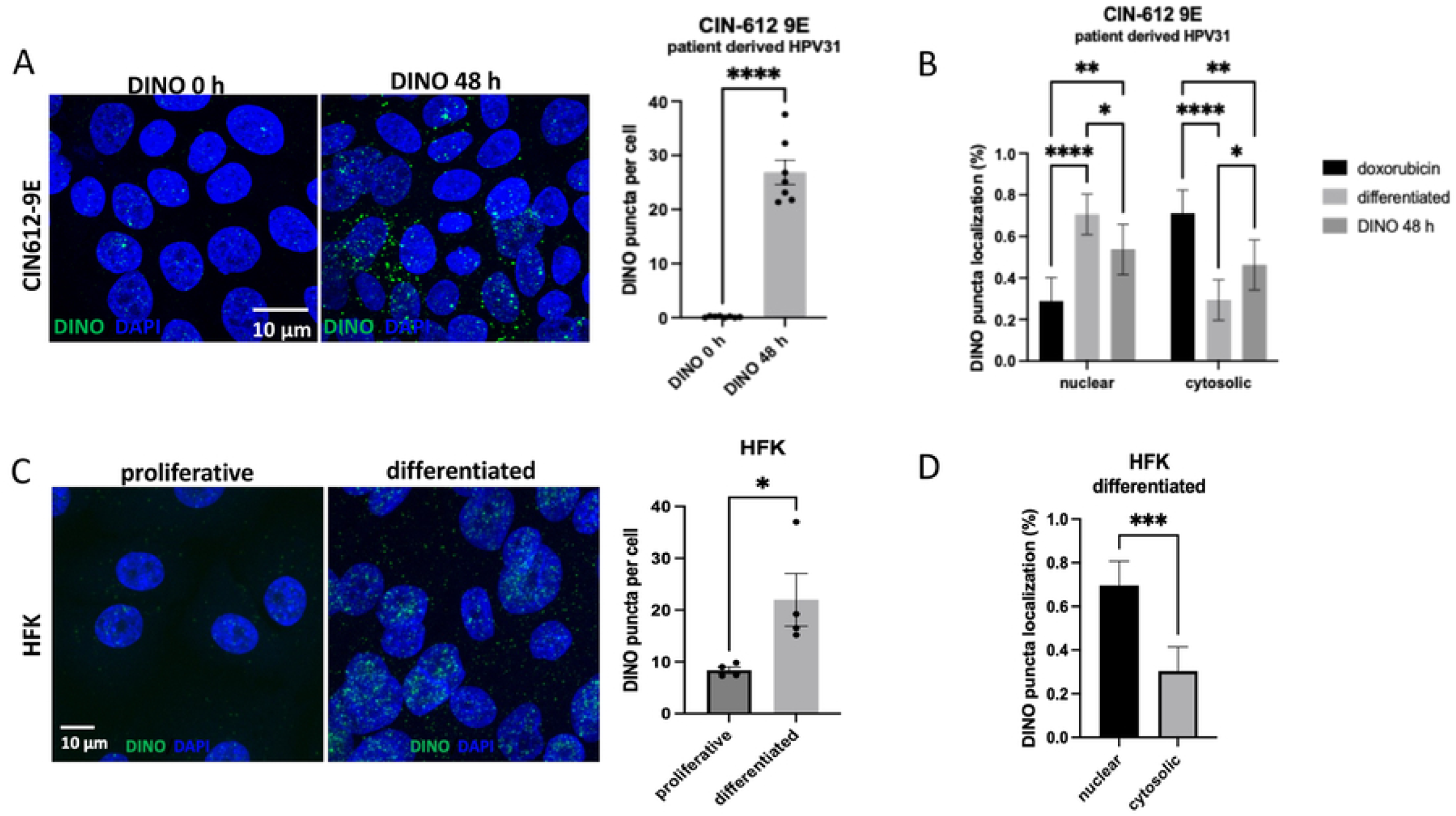
Nuclear DINO transcripts are enriched during differentiation in both HPV-containing cells and primary keratinocytes. (A) RNAscope analysis was conducted in CIN612-9E cells following 48 hours of doxycycline treatment. Quantification of DINO puncta per nucleus indicates an average of ∼25 transcripts per nucleus. Data are presented as mean ± SEM from one biological replicate with two technical replicates (two coverslips); each data point represents one image containing approximately 25 nuclei. The experiment was repeated three times with similar results. Statistical comparisons were performed using an unpaired t-test with Welch’s correction. (B) Quantification of DINO transcript subcellular localization showing the percentage of cytoplasmic versus nuclear signal in doxorubicin-treated, differentiated, and doxycycline-induced DINO-expressing CIN612-9E cells. Data are presented as mean ± SEM from four biological replicates. Statistical comparisons were performed using two-way ANOVA. (C) RNAscope analysis of DINO transcripts in primary human foreskin keratinocytes (HFKs) under proliferative or differentiated conditions. Quantification of DINO puncta per nucleus indicates an average of ∼22 transcripts per nucleus upon differentiation. Data are presented as mean ± SEM from one biological replicate with two technical replicates (two coverslips); each data point represents one image containing approximately 25 nuclei. The experiment was repeated twice with similar results. Statistical comparisons were performed using an unpaired t-test with Welch’s correction. (D) Quantification of DINO transcript subcellular localization in differentiated HFKs, showing the percentage of cytoplasmic versus nuclear signal. Data are presented as mean ± SEM from two biological replicates. An unpaired t-test was used for statistical comparisons.

### Ectopic DINO expression promotes HPV genome replication

Previous studies have shown that HPVs induce breaks in both cellular and viral DNAs, and that breaks found in viral episomes are preferentially repaired, ultimately resulting in genome amplification [30]. Hence, we hypothesized that the enhanced DNA damage signaling associated with DINO expression may trigger HPV genome replication.

Acute DINO expression in W12e cells significantly increased HPV16 genome copy numbers compared to doxycycline-treated GFP expressing control cells, at 48 (2.4 +/- 0.2 fold) and 72 hours (2.3 +/- 1.1 fold) post DINO induction (Figure 5A, left panel) and CIN-612 9E cells at 48 (1.8 +/- 0.9 fold) and 72 (2.2 +/- 1.1 fold) hours post DINO induction (Figure 5B, left panel). To compare these increases to differentiation-induced HPV genome amplification, we determined the increase in HPV genome levels in W12e and CIN612-9E cells in response to Ca^2+^-induced differentiation. In W12e cells, DINO expression resulted in a greater increase in genome copies (∼2 fold) compared to differentiation (∼1.4 fold) (Figure 5A, right panel). In contrast, differentiation contributed more substantially to genome amplification in the CIN-612 9E cell line with ∼8 fold increase in viral DNA compared to the 2-fold induction after DINO expression (Figure 5B, right panel). Since the CIN-612 9E cells amplified viral genomes during differentiation to a greater extent than the W12e cell line, and W12e cells are more prone to undergo HPV genome integration than CIN612-9E cells, we used CIN612-9E cells for further experiments.

**Figure 5.**
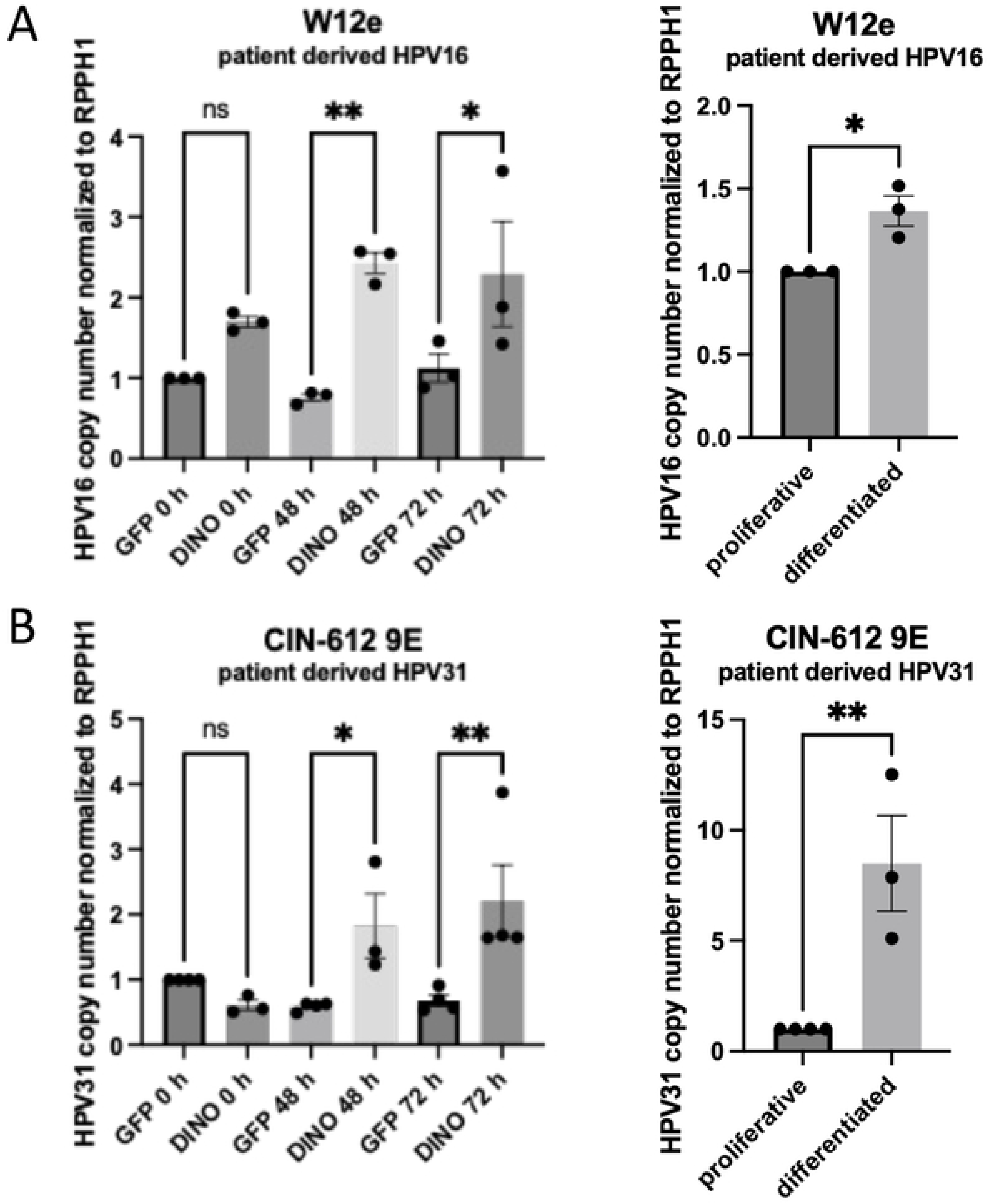
Ectopic DINO expression promotes HPV genome replication. (A) W12 cells transduced with a doxycycline-inducible DINO expression vector were treated with doxycycline and harvested at 0, 48, and 72 hours. HPV16 genome copy number was quantified by qRT-PCR using a standard curve and normalized to the host gene RPPH1. Data are presented as mean ± SEM from three independent experiments and expressed as fold change relative to the 0-hour time point. Statistical comparisons were performed using one-way ANOVA with comparisons between DINO- and GFP-expressing cells at each time point. The panel on the right shows HPV16 copy number in parental proliferative and differentiated W12 cells, with statistical comparison by unpaired t-test. (B) CIN612-9E cells transduced and treated in the same manner were assessed for HPV31 genome copy number by qPCR using a standard curve, normalized to RPPH1, and expressed as fold change relative to 0 hours. Data are presented as mean ± SEM from three independent experiments. Statistical comparisons were performed using one-way ANOVA with comparisons between DINO- and GFP-expressing cells at each time point. The panel on the right shows HPV31 genome copy number in parental proliferative and differentiated CIN612-9E cells, with statistical comparison by unpaired t-test.

### HPV31 viral transcripts are upregulated during keratinocyte differentiation

The HPV31 genome encodes multiple transcripts that are differentially expressed across the viral life cycle (Figure 6A). To establish a baseline for transcript expression in our model system, we measured the levels of the E1, E2, E8^E2, E6*, E7, E1^E4, and L1* mRNAs by qRT-PCR in CIN612-9E cells under proliferative and differentiated conditions. Consistent with the known biology of the HPV life cycle, all viral transcripts were significantly upregulated upon differentiation compared to proliferative conditions (Figure 6B). These findings confirm that our model system faithfully recapitulates the differentiation-dependent transcriptional program of HPV31 and establish a reference for transcript levels against which DINO-induced changes can be compared.

**Figure 6.**
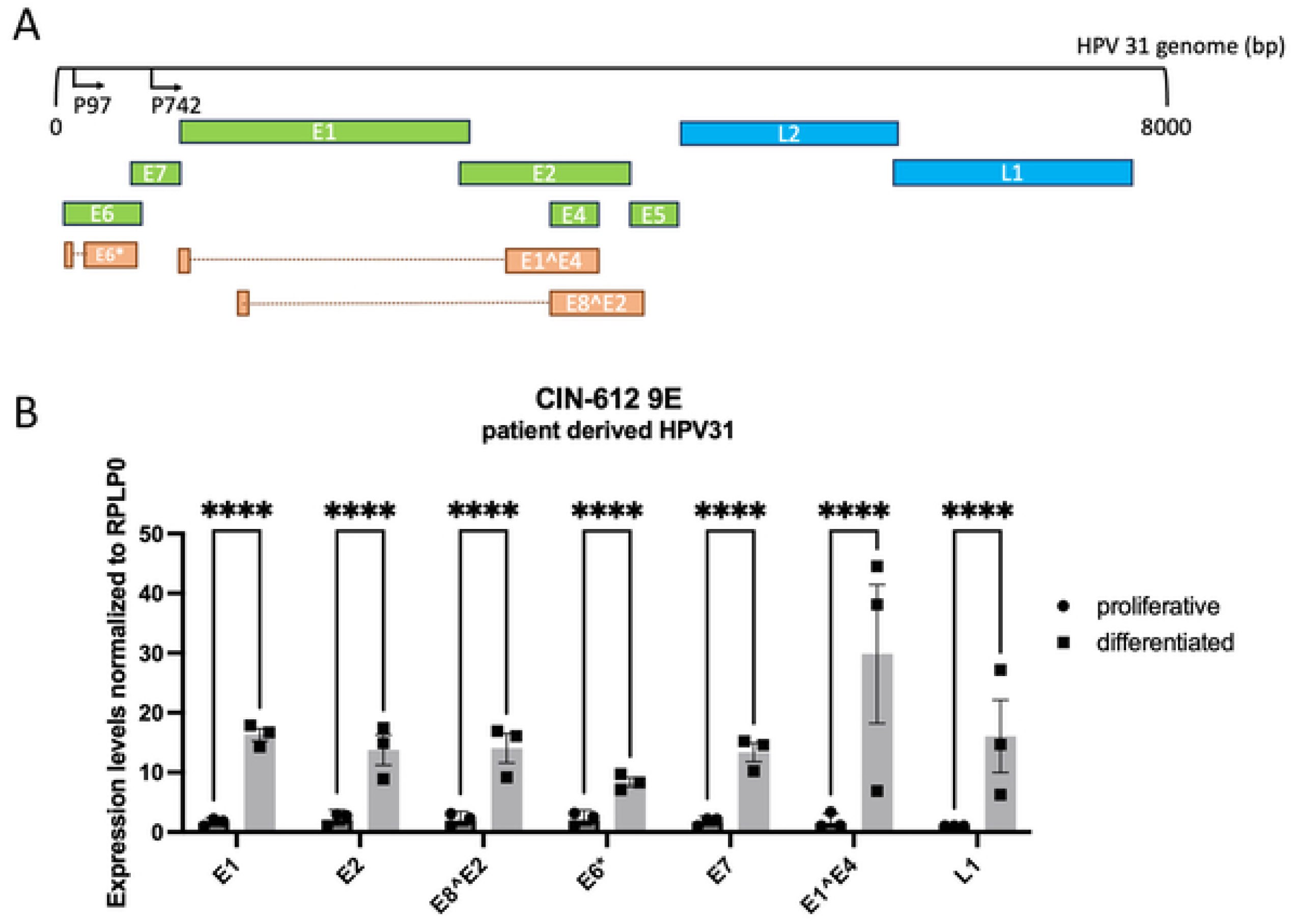
HPV31 viral transcripts are upregulated during keratinocyte differentiation. (A) Schematic of the HPV31 genome depicting the major viral transcripts expressed during the HPV life cycle, including E1, E2, E8^E2, E6*, E7, E1^E4, and L1*. The transcript map was generated based on HPV31 genome annotation available through PAVE (Papillomavirus Episteme) [57, 58]. (B) Proliferative or differentiated CIN612-9E cells were harvested and viral transcript levels were measured by qRT-PCR. Expression of E1, E2, E8^E2, E6*, E7, E1^E4, and L1* were all significantly upregulated during differentiation compared to proliferative conditions. Data are presented as mean ± SEM from three independent experiments, each with two technical replicates. Statistical comparisons were performed using two-way ANOVA with multiple comparisons between proliferative and differentiated conditions.

### Ectopic DINO expression selectively upregulates viral transcripts without triggering keratinocyte differentiation

E1^E4 is one of the most abundantly expressed viral proteins during the productive phase of the HPV life cycle and contributes to viral genome amplification and late gene expression [48, 49]. Some evidence suggests that E1^E4 may also play a role in genome replication in basal cells [50]. Given its established role in promoting productive replication and its expression pattern correlating with increased viral genome copy number, we hypothesized that ectopic DINO expression might increase HPV31 E1^E4 transcript levels. To test this, we measured the effect of acute DINO expression on a panel of viral transcripts representing both early and late stages of the HPV31 transcriptional program.

In CIN612-9E cells, acute DINO expression resulted in a modest yet statistically significant upregulation of E1 and E2 transcripts at 48 hours compared to GFP expressing control cells (Figure 7A). The E1^E4 transcript levels were more highly increased than the E1 and E2 transcripts at 48 hours (Figure 7A), although these levels did not reach those observed during differentiation (Figure 6B). In contrast, E8^E2, E6*, E7, and the late capsid transcript L1* remained unchanged upon DINO expression (Figure 7A), indicating that DINO selectively increases the abundance of a subset of viral transcripts rather than broadly activating the HPV31 transcriptional program as is observed during differentiation.

**Figure 7.**
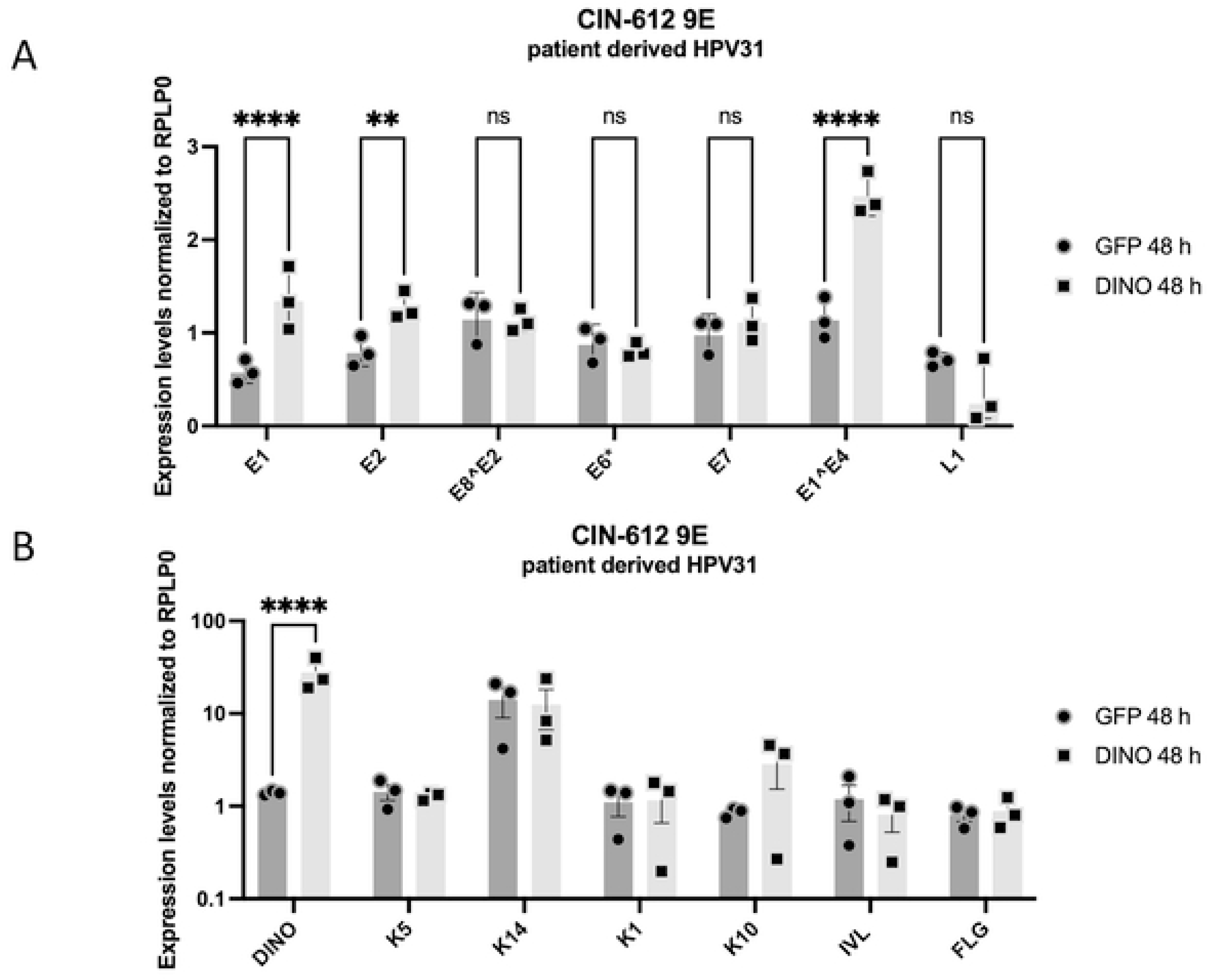
Ectopic DINO expression selectively upregulates viral transcripts without triggering keratinocyte differentiation. (A) CIN612-9E cells expressing doxycycline-inducible GFP or DINO were harvested at 0- and 48-hours post-doxycycline treatment. qRT-PCR was performed to measure expression levels of E1, E2, E8^E2, E6*, E7, E1^E4, and L1*. DINO expression resulted in the modest upregulation of E1 and E2, and a more pronounced increase in E1^E4 transcript levels at 48 hours compared to GFP controls, although E1^E4 induction did not reach the levels observed during differentiation. E8^E2, E6*, E7, and L1* transcript levels remained unchanged. Data are presented as mean ± SEM from three independent experiments, each with two technical replicates. Statistical comparisons were performed using two-way ANOVA with multiple comparisons between DINO- and GFP-expressing cells at each time point. 0-hour timepoints were included in all statistical analyses but omitted from graphical representation for clarity. (B) qRT-PCR was performed on the same samples to measure expression of DINO and a panel of keratinocyte differentiation markers representing distinct stages of differentiation: KRT5 and KRT14 (basal/proliferative), KRT1 and KRT10 (early differentiation), and IVL and FLG (late differentiation). Data are presented as mean ± SEM from three independent experiments, each with two technical replicates. Statistical comparisons were performed using two-way ANOVA with multiple comparisons between DINO- and GFP-expressing cells at each time point. 0-hour timepoints were included in all statistical analyses but omitted from graphical representation for clarity.

Since endogenous DINO levels increase during the differentiation of CIN612-9E cells (Fig. 2B) we sought to investigate whether DINO may promote viral replication indirectly by driving cellular differentiation. To address this we acutely expressed DINO or GFP as a negative control, in CIN612-9E cells and measured expression by qRT-PCR of a panel of canonical early and late differentiation markers including keratins K5 and K14 that are expressed in undifferentiated cells, keratins K1 and K10 that are early keratinocyte differentiation markers and involucrin (IVL), filaggrin (FLG) which are late differentiation markers, [51]. Acute DINO expression did not significantly alter the expression of any of these differentiation-associated genes compared to GFP expressing controls (Figure 7B), demonstrating that the selective upregulation of viral transcripts by DINO occurs independently of keratinocyte differentiation.

### DINO expression causes an increase in HPV replication foci

Differentiation-dependent HPV genome amplification occurs at distinct replication foci [52]. Next, we determined whether the observed increase in HPV genome copy number in response to doxycycline-mediated, acute DINO expression also caused an increase in HPV replication foci. We used fluorescence in situ hybridization (FISH) to directly visualize HPV31 genomes (Figure 8A). As expected, CIN612-9E cells exhibited sparse replication foci when grown under undifferentiated conditions. The proportion of HPV31 FISH foci containing CIN612-9E cells markedly increased when the cells were induced to differentiate by treatment with 1.5 mM CaCl_2_ for 96 hours.

**Figure 8.**
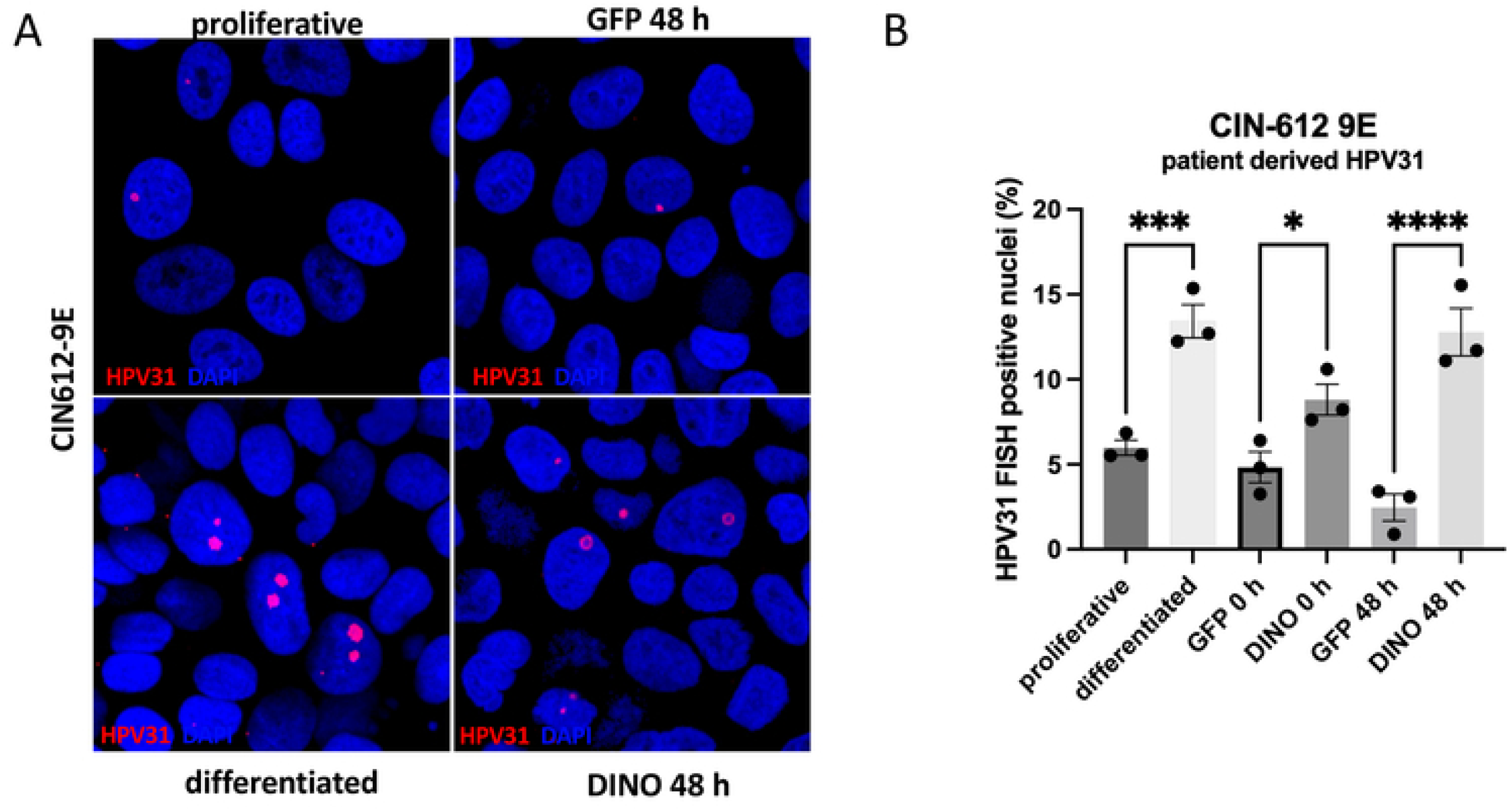
Ectopic DINO expression causes an increase in HPV replication foci. (A) Representative fluorescence microscopy images of CIN612-9E cells under proliferative, differentiated, or doxycycline-induced conditions (48-hour induction of GFP or DINO). Cells were hybridized with an HPV31-specific FISH probe to visualize viral replication foci and counterstained with DAPI to mark nuclei.(B) Quantification of HPV31 replication foci based on blinded scoring of approximately 300 nuclei per condition per replicate (three replicates). The percentage of nuclei containing detectable HPV replication foci is shown for proliferative, differentiated, GFP-, and DINO-expressing cells at 0 and 48 hours. Differentiated cells exhibited a robust increase in replication foci, while ectopic DINO expression led to a measurable increase at 48 hours post-induction, with a modest elevation also observed at 0 hours compared to GFP controls. Data are presented as mean ± SEM. Statistical comparisons were performed using one-way ANOVA with multiple comparisons between DINO- and GFP-expressing cells at each time point, as well as between proliferative and differentiated conditions.

Acute, doxycycline-mediated DINO expression for 48 hours also led to a significant increase in HPV31 replication foci, reaching levels comparable to those observed during differentiation. In contrast, acute doxycycline-mediated GFP expression for 48 hours did not cause an increase in HPV replication foci. Quantification of FISH-positive nuclei (Figure 8B) confirmed that similar to Ca^2+^-mediated differentiation, DINO induction significantly elevated the percentage of cells with detectable replication foci. These results show that the increase in HPV genome copy number in response to DINO expression is associated with the formation of viral replication foci.

## Discussion

In this study, we show that endogenous DINO levels increase during keratinocyte differentiation in both HPV negative and positive keratinocytes. We further demonstrate that DINO enhances HPV replication independently of canonical differentiation signals, as ectopic DINO expression did not induce keratinocyte differentiation markers. Notably, DINO’s effects on HPV replication also appear to be independent of p53, as ectopic DINO expression did not result in p53 stabilization or upregulation of the canonical p53 target *CDKN1A*. This is particularly significant given that DINO was originally characterized as a p53-dependent lncRNA [34] and suggests that DINO can engage alternative regulatory pathways in the context of HPV-positive keratinocytes. Ectopic DINO expression significantly increased HPV genome copy number, the formation of viral replication foci, and levels of the viral transcripts E1, E2, and E1^E4. E1^E4 plays a critical role in genome amplification and maintenance during the viral life cycle, particularly in the context of keratinocyte differentiation [50]. The modest upregulation of E1 and E2 is also noteworthy, as these encode the viral helicase and origin-binding protein respectively, which are essential for viral genome replication, and their increased expression may contribute to the observed increase in genome copy number.

Importantly, the lack of induction of the L1* late viral transcript suggests that DINO does not fully activate the late viral transcriptional program, and its effects appear confined to transcripts associated with early viral genome replication functions. Whether the observed increase in viral transcripts results from direct DINO-mediated transcriptional activation or is a consequence of the observed enhanced HPV genome replication remains to be elucidated. However, the selective upregulation of E1, E2, and E1^E4 without a corresponding increase in other viral transcripts argues against a simple copy number-driven effect, as global upregulation of viral transcripts is observed during differentiation when genome amplification occurs. The selective increase in E1, E2, and E1^E4 without corresponding changes in E6 and E7, which are present on the same polycistronic transcripts and would be expected to increase proportionally if transcription initiation were enhanced, raises the possibility that DINO influences viral transcript stability or processing rather than transcription initiation per se.

The doxycycline-inducible system employed in this study increased DINO expression to levels comparable to those observed in response to induction of DNA damage by doxorubicin treatment or by differentiation, ensuring physiological relevance. However, we did not test RNA stability, and it is possible that doxorubicin or differentiation may not only induce DINO transcription but could also affect transcript stability. To understand the subcellular localization patterns of DINO transcripts under different conditions, we employed RNA scope in CIN612-9E cells.

Endogenous DINO expression was assessed in control cells, doxorubicin-treated cells, and in cells undergoing differentiation. Both doxorubicin treatment and differentiation increased DINO expression compared to control conditions, as evidenced by the elevated number of RNA scope puncta per cell. However, differentiation was associated with a pronounced nuclear enrichment of DINO transcripts regardless of HPV presence, whereas doxorubicin treatment resulted in a predominantly cytoplasmic localization of DINO. DINO has been reported to occupy p53-bound regulatory regions on chromatin [34], consistent with a possible nuclear regulatory role, but a specific cytoplasmic function for DINO remains unknown. These contrasting patterns suggest distinct regulatory mechanisms driving DINO transcript localization during epithelial differentiation and DNA damage and it is unknown whether the nuclear DINO pool is connected to its reported p53 related biological activities.

To evaluate whether ectopic DINO expression mirrors the localization patterns observed under DNA damage or differentiation conditions, we analyzed CIN612-9E cells after 48 hours of doxycycline-induced DINO expression by RNA scope. The number of DINO puncta in doxycycline-induced cells closely resembled that of doxorubicin-treated cells. However, the subcellular localization pattern was intermediate, with transcripts almost equally distributed between the cytoplasm and nucleus. This localization pattern suggests that acute DINO expression partially mirrors the regulatory dynamics observed during DNA damage and differentiation. These findings highlight the utility of the doxycycline-induced system in modeling aspects of DINO expression, while also underscoring the complexity of endogenous DINO regulation under distinct cellular conditions.

DINO expression leads to an increased number of γH2AX foci per cell, although we cannot conclusively attribute this to an elevation in double-strand DNA break events. DINO may instead interfere with the repair of double-strand DNA breaks or hinder replication fork progression, resulting in stalled or collapsed replication forks and persistent γH2AX signaling, or may influence replication-associated chromatin in ways that promote a DNA damage-like environment favorable for HPV replication. Future studies using high-resolution imaging and biochemical analyses will be critical to dissect the molecular interactions between DINO, DNA repair machinery, and viral replication foci.

Together, these observations suggest that stress-responsive lncRNAs such as DINO may contribute to creating an intracellular environment conducive to HPV genome amplification. In proliferating keratinocytes, DINO expression is minimal; however, DINO levels rapidly increase in response to cellular stressors such as DNA damage or differentiation, independently of canonical differentiation signals. HPV may therefore exploit stress-responsive lncRNAs like DINO to initiate or enhance replication during times of cellular upheaval, a strategy observed in other viruses that co-opt host lncRNAs to enhance replication or evade immune defenses [53, 54].

However, our results also indicate that while DINO contributes to genome amplification, it is not sufficient to drive the complete HPV vegetative life cycle. The ability of ectopic DINO expression to increase the number of viral replication foci and modestly upregulate E1, E2, and E1^E4 transcripts and genome copy numbers suggests that DINO plays a significant role in establishing a replication-competent state. The increase in E1 and E1^E4 transcripts is consistent with a role for DINO in promoting viral genome replication, as E1 encodes the viral helicase required for origin unwinding and replication initiation, and E1^E4 has been shown to localize to and support viral replication compartments [48, 55, 56] .The failure to induce L1* transcripts or robust genome amplification comparable to vegetative DNA amplification indicates that additional differentiation-dependent mechanisms are required for progeny virus synthesis, such as differentiation-induced chromatin remodeling, accumulation of viral replication factors, or other host factors yet to be identified.

This study opens several avenues for further research into the role of DINO in HPV biology. Although we show that DINO levels increase during keratinocyte differentiation and that ectopic DINO expression is sufficient to induce HPV genome replication in the absence of differentiation, it is not clear whether differentiation-induced DINO induction is necessary for HPV genome amplification Although loss-of-function approaches were pursued, siRNA and shRNA-mediated knockdown of DINO during differentiation achieved limited efficiency, precluding definitive conclusions regarding the impact of DINO depletion on differentiation-dependent HPV genome amplification. Future studies employing more complete depletion strategies will be critical to determine whether DINO is necessary to promote the productive phase of the HPV life cycle. We also have limited insights regarding the mechanism by which DINO may drive HPV genome amplification. Our results suggests that the mechanism is unrelated to p53, and future analyses of the transcriptional responses of DINO expression in CIN612-9E cells may provide insights. Future work will also focus on investigating whether DINO directly interacts with viral replication factors, such as E1 or E2, or with host proteins involved in replication fork stability and repair will provide insights into its molecular mechanism of action. Additionally, the mechanisms driving DINO’s selective impact on certain viral transcripts, without affecting the expression of late transcripts such as L1, remain unclear and warrant further exploration. Understanding these selective effects could reveal critical regulatory checkpoints in HPV replication.

## Acknowledgments

We thank Drs Paul Lambert (University of Wisconsin, Madison) and Laimionis Laimins for providing the W12e and CIN612-9E cells, respectively, Dr. Ralph Isberg, Claire Moore, Phil Hinds, and Marta Gaglia and members of the Munger Lab, particularly Wendelin Marmol for stimulating discussions and Miranda Grace for assistance with western blot experiments. This work was supported by the National Institutes of Health, grant number R01 AI170633 (K.M.), and National Institutes of Health, and T32GM139772 (W.A.). AAM is supported by the Intramural Research Program of the National Institute of Allergy and Infectious Diseases, of the National Institutes of Health (NIH). The contributions of the NIH author are considered Works of the United States Government. The findings and conclusions presented in this paper are those of the authors and do not necessarily reflect the views of the NIH or the U.S. Department of Health and Human Services.

